# Using RNA-seq for genomic scaffold placement, correcting assemblies, and genetic map creation in a common *Brassica rapa* mapping population

**DOI:** 10.1101/076745

**Authors:** RJ Cody Markelz, Michael F Covington, Marcus T Brock, Upendra K Devisetty, Daniel J Kliebenstein, Cynthia Weinig, Julin N Maloof

**Affiliations:** University of California at Davis, Department of Plant Biology, Davis, CA 95616; University of California at Davis, Department of Plant Sciences, Davis, CA 95616; University of Wyoming, Department of Botany, Laramie, WY, 82072; Amaryllis Nucleics, Inc. Berkeley, CA, 94710

**Keywords:** RNA-seq, genetic map, *Brassica rapa*, genome assembly correction

## Abstract

*Brassica rapa* is a model species for agronomic, ecological, evolutionary and translational studies. Here we describe high-density SNP discovery and genetic map construction for a *Brassica rapa* recombinant inbred line (RIL) population derived from field collected RNA-seq data. This high-density genotype data enables the detection and correction of putative genome mis-assemblies and accurate assignment of scaffold sequences to their likely genomic locations. These assembly improvements represent 7.1-8.0% of the annotated *Brassica rapa* genome. We demonstrate how using this new resource leads to a significant improvement for QTL analysis over the current low-density genetic map. Improvements are achieved by the increased mapping resolution and by having known genomic coordinates to anchor the markers for candidate gene discovery. These new molecular resources and improvements in the genome annotation will benefit the *Brassicaceae* genomics community and may help guide other communities in finetuning genome annotations.

## INTRODUCTION

The *Brassica* genus is important for human diets throughout Asia, providing micronutrients and up to 12% of oil calories and a wide diversity of agricultural products (Dixon, 2007; Wang *et al.*, 2011b). Within this genus genome sequences have recently been published for *Brassica napus*, *Brassica rapa*, and *Brassica oleracea*(Chalhoub *et al.*, 2014; Liu *et al.*, 2014; Wang *et al.*, 2011b)*. Brassica rapa*is a physiologically and morphologically diverse diploid species that has 87% gene exon similarity to the model plant *Arabidopsis thaliana*(Cheng *et al.* 2013). This makes *Brassica rapa*an excellent species for comparing and translating knowledge of biological processes from Arabidopsis to a crop species. For example, homologous Arabidopsis gene information has been used to infer the action of *B. rapa*genes in glucosinolate metabolism (Li and Quiros, 2001; Wang *et al.*, 2011a), flowering time, leaf development (Baker *et al.*, 2015), and seed yield (Brock *et al.*, 2010; Dechaine *et al.*, 2014). All of these important traits contribute to our understanding of plant growth in agricultural settings and the underlying genetic understanding of these traits is made possible by a reference genome sequence (Wang *et al.*, 2011b), gene annotation information (Cheng *et al.*, 2013; Devisetty *et al.*, 2014), and genetic mapping populations (e.g. Iniguez-Luy *et al.* 2009).

The annotated *Brassica rapa*genome assembly is 283.8 Mb spread over 10 chromosomes A01-10 (Wang *et al.* 2011b). Although the current genome is diploid, there are three ancient subgenomes derived from genome duplication events. These subgenomes are designated as least fractionated (LF), most fractionated one (MF1), and most fractionated two (MF2) corresponding to the fraction of gene loss in each subgenome (Cheng et al., 2012; Wang et al., 2011b). These three subgenomes share many paralogous genes and contiguous regions complicating genome assembly. This has prevented about 10.8% of the gene-containing genomic scaffolds in version

1.5 of the genome (http://brassicadb.org/brad/index.php) from being assigned to chromosomes. The lack of chromosomal assignment is largely because these scaffolds have no molecular markers that would have enabled their placement on the genetic map. This suggests that identifying more markers can help to make the *B. rapa*genome assembly more comprehensive (Wang *et al.*, 2011b).

For this study we utilized an existing RIL population of *Brassica rapa*that has been used extensively for QTL mapping of physiological, developmental, and evolutionarily important traits (Bra-IRRI; Baker et al., 2015; Brock et al., 2010; Dechaine et al., 2007, 2014; Edwards et al., 2011; Iniguez-Luy et al., 2009; Lou et al., 2011, 2012). Recently, we completed deep RNA-sequencing of the parents of the Bra-IRRI population providing a large SNP set and improved gene annotation information (Devisetty *et al.*, 2014). Using a new set of RNA-seq data collected on the entire population, we extend these SNP discovery methods to 124 genotypes in the population for placing scaffolds, correcting assemblies, and the creation of a saturated genetic map.

## MATERIALS AND METHODS

### Plant Growth and Tissue Collection

The field site for plant growth was located at the University of Wyoming Agricultural Experimental Station in Laramie, Wyoming, USA. This study focused on 124 RILs and the two parental genotypes (R500 and IMB211) of the *Brassica rapa*IRRI population. Individual replicates of each RIL were sown into peat pots filled with field soil and topped with 1 cm LP5 potting soil (Sun Gro Horticulture, Agawam, MA, USA). Seeds were planted in the first week of June 2011, and pots were transplanted to the field 2.5 weeks later following established protocols (Dechaine *et al.*, 2014). One biological replicate of each genotype was planted into each of 5 fully randomized blocks. After plants were established in the field for three weeks, apical meristem tissue was collected from individual replicate plants into 1.5 mL Eppendorf tubes, immediately flash frozen in liquid nitrogen, and stored at -80 ºC until RNA-Seq library preparation.

### NA-Seq library preparation and sequencing

RNA-Seq libraries were prepared using a high-throughput Illumina RNA-Seq library extraction protocol (Kumar *et al.*, 2012). The enriched libraries were then quantified on an Analyst Plate Reader (LJL Biosystems) using SYBR Green I reagent (Invitrogen). Once the concentration of libraries was determined, a single pool of all the RNA-Seq libraries within each block was made. The pooled libraries were run on a Bioanalyzer (Agilent, SantaClara) to determine the average product size for each pool. Each pool was adjusted to a final concentration of 20 nM and sequenced on 7 lanes on Illumina Hi-Seq 2000 flow cell as 50-bp single end reads. Any failed samples from the 5 blocks were run on 2 additional lanes.

### NA-Seq Read Processing

Pre-processing and mapping of Illumina RNA-Seq raw reads was done as described in detail in Devisetty *et al.* 2014 with one exception. The raw reads were quality filtered with FASTX tool kit’s (http://hannonlab.cshl.edu/fastx_toolkit/) fastq_quality_filter with parameters [-q 20, -p 95]. The qualified de-multiplexed reads were then mapped to *B. rapa*reference genome (Chiifu version 1.5) using BWA v0.6.1-r104 Li and Durbin, 2009 with parameters [bwa_n 0.04] and the unmapped reads were in turn mapped with TopHat with parameters [splice-mismatches 1, max-multihits 1, segment-length 22, butterfly-search, max-intron-length 5000, library-type frunstranded]. Finally, mapped reads from both BWA and TopHat were combined for genotyping purposes.

### Population-based Polymorphism Identification

Variant Call Format (VCF) files were generated for each of five replicate blocks of samples using samtools and bcftools. These tools were run as 'samtools mpileup -E -u -f Brapa_sequence_v1.5.fa [all alignment files for the current block] | bcftools view -bvcg - | vcfutils.pl varFilter'.

The VCF files were summarized using 'summarize-vcf.pl' Perl script (https://github.com/mfcovington/snps-from-rils). For each block of replicates, this script (run using the parameters: '--observed_cutoff 0.3 --af1_min 0.3') ignores INDELs and variant positions with more than two alleles, ignores variants with site allele frequency (AF1) values too far from 0.5 (>= 0.7 or <= 0.3), and ignores variants with missing information in 30% or more of the population. For variants that passed these filters, the numbers of reads matching the reference and the number of alternate allele reads were recorded in a VCF summary file.

These VCF summary files from the different replicate blocks were merged using the 'merge-vcf-summaries.pl' (https://github.com/mfcovington/snps-from-rils) Perl script. Using the default parameters ('--replicate_count_min 2 --ratio_min 0.9'), this script merges the information in the VCF summaries and records a putative SNP as an actual SNP if the variant is identified as a SNP in at least 2 replicate blocks and if the proportion of reads matching the major allele is at least 0.9. This was done on a RIL by RIL basis.

### Genotyping, Plotting, and Identification of Genotype Bins

The Perl script 'extract+genotype_pileups.pl' (https://github.com/mfcovington/detect-boundaries) was used with the '--no_nr' parameter to extract genotype information from the RNA-seq alignments at each SNP location for each member of the RIL population. The resulting genotype files were used to detect and remove SNPs with excessive noise.

Due to the crossing scheme used to create the RIL population, each individual is expected to be nearly homozygous for one parent or the other. The 'filter-noisy-SNPs.pl' (https://github.com/mfcovington/noise-reduction-for-snps-from-pop) Perl script performs noise-reduction for SNPs derived from such a population. It does this by identifying and ignoring positions that have an over-representation of heterozygosity in individual lines across the entire population. Using the default parameters ('--cov_min 3 --homo_ratio_min 0.9 -- sample_ratio_min 0.9'), SNPs were discarded as noisy if more than 10% of the lines in the population showed evidence of heterozygosity as defined by a line having at least 3 reads per SNP position with a major allele with a ratio less than 0.9.

After noise-reduction, the 'extract+genotype_pileups.pl' (https://github.com/mfcovington/SNPTools) Perl script was re-run without the '--no_nr' parameter for each RIL. The resulting genotype files were used to create genotype plots using the 'genoplot_by_id.pl' Perl script (https://github.com/mfcovington/SNPTools) and to define genotype bins for the individual RILs.

The 'filter-snps.pl' Perl script (https://github.com/mfcovington/detect-boundaries) was used to identify regions of adjacent SNPs with alleles from the same genotype. Using the default parameters ('--min_cov 10 --min_momentum 10 --min_ratio 0.9 --offset_het 0.2'), it detects boundaries between genotype bins when a sliding window of at least 10 SNPs. Within each sliding window a depth of at least 10 reads each exhibit major allelic ratios of at least 0.9. The major allele represents the opposite genotype from the previous bin (or exhibit major allelic ratios less than 0.7 for transitions from regions of homozygosity to those of heterozygosity). For each member of the RIL population, this script generates one file with boundary between genotype bins.

The 'fine-tune-boundaries.pl' Perl script (https://github.com/mfcovington/detect-boundaries) is an automated tool for rapid, fine-scale human curation of boundaries between genotype bins that we used for the RIL population. As described in Devisetty et al. 2014, "This command-line tool displays color-coded genotype data together with the currently-defined bin boundaries. Using shortcut keys, the operator can quickly and easily approve or fine-tune a boundary (at which point, the next boundary is instantly displayed for approval)."

The 'merge-boundaries.pl' Perl script (https://github.com/mfcovington/detect-boundaries) was used to merge all of the boundaries in the collection of the boundaries files that were generated by 'filter-snps.pl' and 'fine-tune-boundaries.pl'. A comprehensive list of bins and their locations resulting from the merge are written to a file: bins.tsv. The script also prints the boundary and bin stats (count, min size, max size, and mean size) to the screen to allow visual analysis of the resulting file. This information was used for human curation of the boundaries.

The 'get-genotypes-for-bins.pl' Perl script (https://github.com/mfcovington/detect-boundaries) was used to convert the comprehensive bins file and all the individual boundaries files into a summary of bins and their locations across the genome and their genotypes across the entire RIL population (Supplemental Table 1).

Composite genotype plots (Figure 3) were created using the 'plot-composite-map.R' R script (https://github.com/mfcovington/detect-boundaries).

### Validating and Reassigning Genomic Scaffolds

Using the genotypic value for each genotype bin across the RILs, we calculated the asymmetric binary distance between all central SNP pairs using the dist(method = “binary”) function in R. The pairwise correlation matrix was then ordered by maximal correlations to place the map in a linear order and compared to the predicted bin order based on version 1.5 of the *Brassica rapa*genome. Comparisons between v1.5 of the genome and binary distance plots were manually inspected to ensure proper placement or reassignment.

### Genetic Map Construction

Because each genotype bin across the RILs represents each observed recombination breakpoint in the population, we used one SNP per genotype bin to create a saturated genetic map. Aside from the possibility of rare, unobserved double cross over events, the mapping resolution in this population is no longer limited by the number of SNPs but instead by recombination events. The genetic map was constructed using the chromosomal position of each of the SNPs as a starting point for marker ordering along the chromosomes. Each chromosome was treated as a large linkage group and each SNP was tested for linkage disequilibrium with all other SNPs using the R/QTL package (Broman *et al.*, 2003) in the R statistical environment (R Core Team, 2015). Larger gaps in RNA-seq information corresponding to low gene density centromeric regions were problematic when ordering markers using the ripple() R/QTL function (Broman *et al.*, 2003). In chromosomes A08 and A09, after local marker order was established we used the physical position of the SNPs to connect the two arms in the correct orientation.

### QTL Comparisons

To test how increased marker coverage affected QTL mapping and identification for physiological traits, we remapped two traits from (Brock *et al.*, 2010) that had been mapped using the previous genetic map (Iniguez-Luy *et al.*, 2009). We used R/QTL (v1.39-5) to compare mapping results derived from the previous and updated genetic maps using the Brock *et al.* (2010) flowering time phenotype data. Specifically we used the cim() function with three marker covariates and determined LOD significance cutoffs after 1,000 permutations.

## RESULTS AND DISCUSSION

### 500 vs. IMB211 polymorphism identification

We performed deep RNA sequencing of 124 individuals of a RIL population derived from a cross between the *Brassica rapa*accessions R500 and IMB211 (Iniguez-Luy *et al.*, 2009). We sequenced five replicates of each RIL and mapped 5.26 million reads mapped per RIL. We had previously identified SNPs and INDELs between R500 and IMB211, the parents of the population (Devisetty *et al.* 2014) using v1.2 of the *Brassica rapa*genome. This set of R500 vs. IMB211 polymorphisms was used to genotype each member of the RIL population individually. The crossing scheme used to create the RIL population should create homozygous regions of contiguous R500 alleles alternating with homozygous regions of contiguous IMB211 alleles in the different RILs. However, when using the R500 vs. IMB211 polymorphism set to genotype the RILs there were multiple regions where R500 and IMB211 alleles were randomly interspersed. This suggested that the RIL population might be derived from a different parent or parents than those that we had sequenced.

To test this hypothesis, we merged the sequence data from all RILs and then genotyped the merged dataset using SNPs identified by the IMB211 vs R500 comparison (Figure 1A). The merged dataset gave us a much better view of segregation of putative parental SNPs in this population. Given the relatively large size of the population and the expected recombination frequency and distribution, polymorphisms identified in the true RIL parents should be segregating with approximately equal allelic frequency in this merged data set (black dots in Figure 1A). Most genomic regions did display this expected distribution; however, there were several large regions that were not segregating, but instead were monomorphic for one of the putative parents of the population (indicated as orange or blue dots in Figure 1A). In other words, SNPs identified as polymorphic between the R500 and IMB211 strains are not segregating in the RILs. Nearly all of these monomorphic regions matched R500 alleles, consistent with the idea that the IMB211 seed strain is not the true parent of the RIL population.

**Figure 1.**
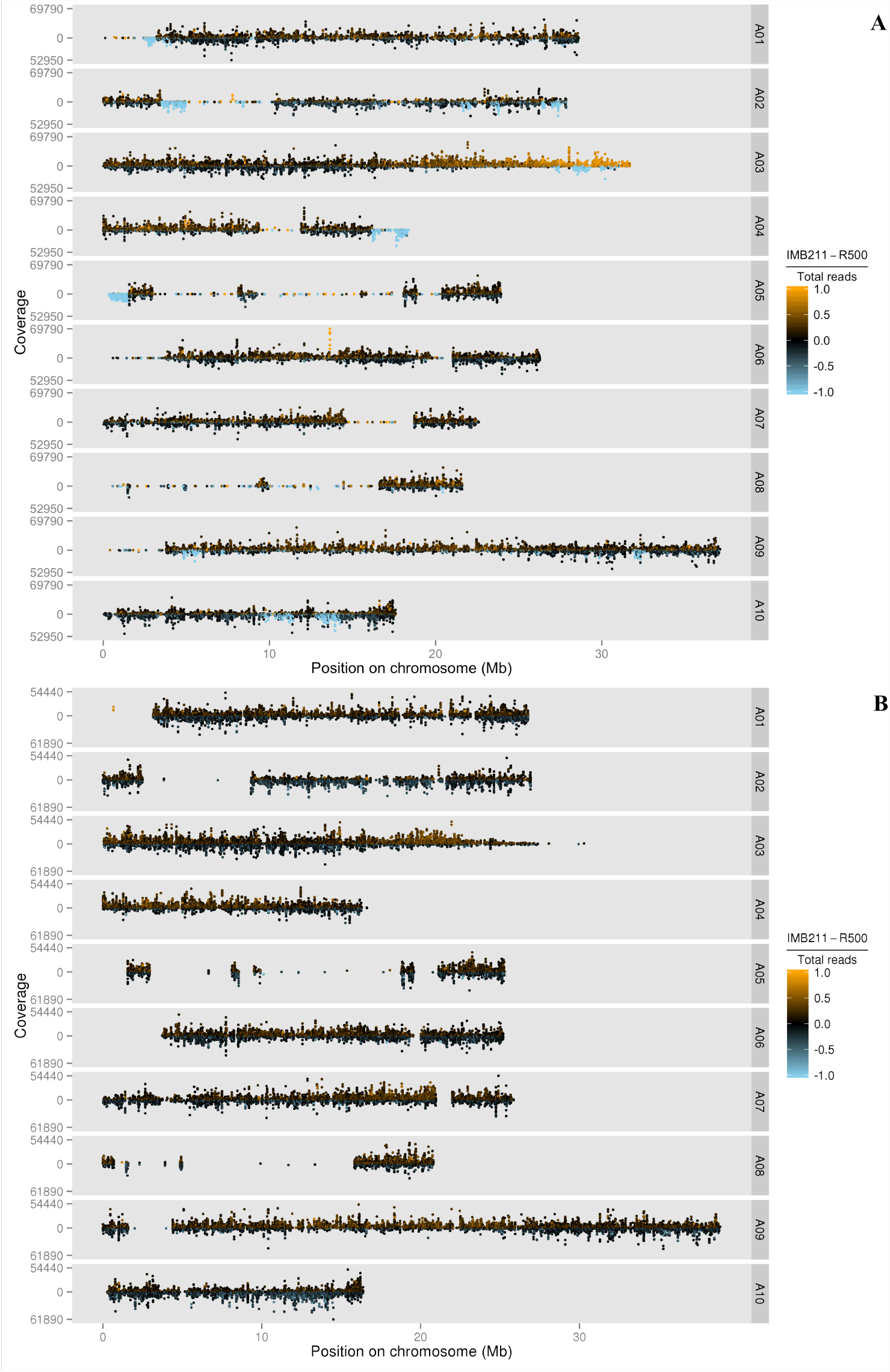
Plot of merged data from all RILs genotyped using the parent-based SNP set (A) and the population based SNP set (B). Each of the *B. rapa*ten chromosomes are displayed (A01-A10) with counts coverage of each SNP at each physical position on the chromosome in megabases (Mb). The color indicates the relative ratio of coverage between R500 and IMB211 for every SNP. Black is equal coverage, orange is more IMB211 and blue is more R500.

The primary exception to the expected, Mendelian parental allele frequency in the RILs is on the bottom of chromosome A03, where there is a gradual transition from equal R500:IMB211 allelic frequency to nearly all IMB211. The A03 pattern is consistent with segregation distortion within the population possibly caused by the centromere being located at that end of chromosome A03 (Cheng *et al.* 2013). With the lower recombination frequencies commonly observed near centromeres in plants (Harushima *et al.*, 1998; Haupt *et al.*, 2001; Sherman and Stack, 1995), there could be a meiotic drive allele or a local inversion in this region causing the segregation distortion and this effect could be enhanced by the proximity to the centromere. There is evidence for each of these mechanisms occurring across a wide range of plant species (Buckler *et al.*, 1999; Fang *et al.*, 2012; Lowry and Willis, 2010).

### Population-based SNP discovery

Due to the uncertainty surrounding the identity of the IMB211 parent of the RIL population, we switched to a population-based approach for SNP discovery. This new strategy involved identifying variants within the RIL population and using the R500 data to assign parental origin for each SNP. Using this approach, we identified 146,027 SNPs across *B. rapa*'s ten chromosomes (Table 1, Supplemental Table 1). These population-based SNPs segregate at the expected allele frequencies of approximately 50/50 throughout the entire genome except at the previously noted end of A03 (Figure 1B). Over 80% of the genome is within 100 kb of a SNP; however, there are several regions with few or no SNPs. There are two primary reasons for these SNP-free regions. Most are likely gene-poor regions or regions of genes with insufficient expression under our experimental conditions (e.g., growth conditions, age, tissue, genotypes). We also found a few regions where there are significant numbers of expressed genes, but no SNPs between members of the RIL population. These regions primarily correspond to the non-variant regions of Figure 1A and, therefore, likely represent regions that are very similar between the seed stocks used to generate this RIL population.

**Table 1.**
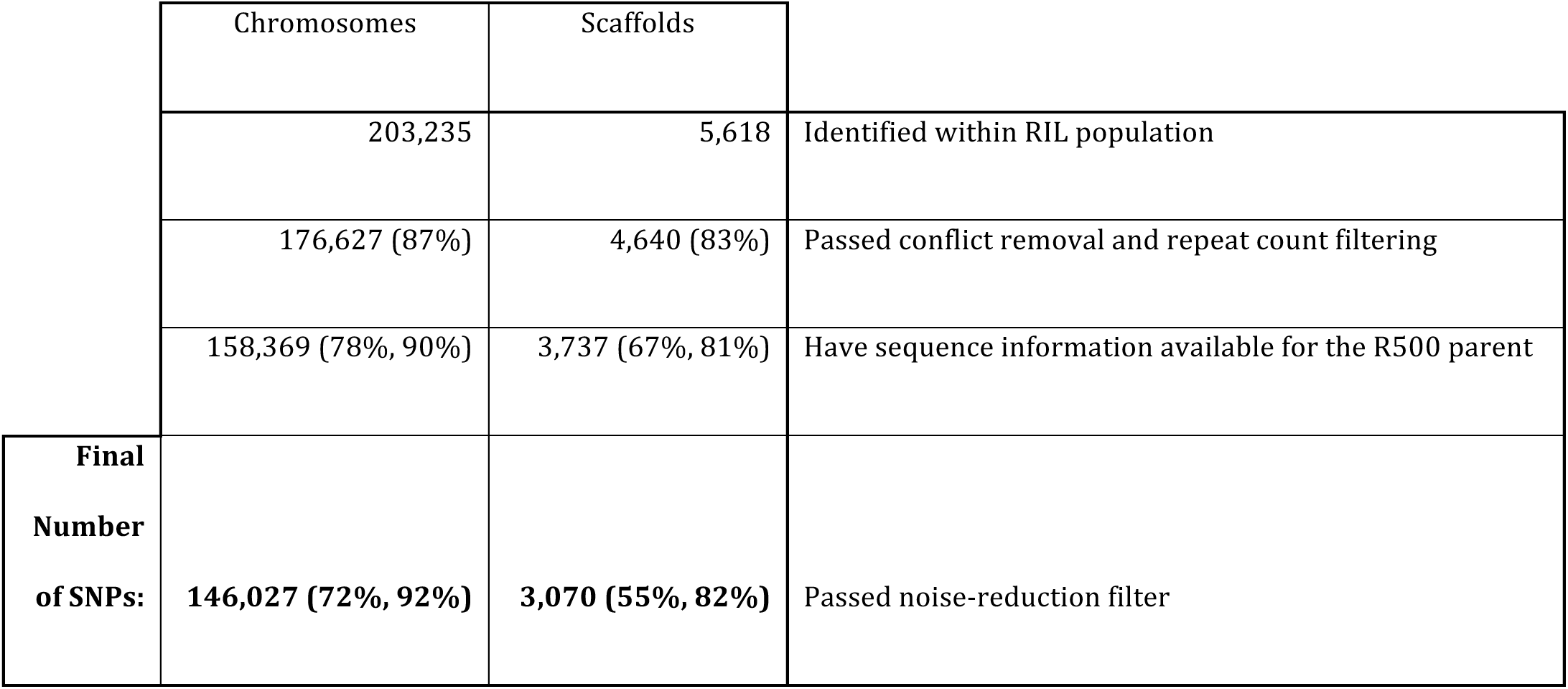
SNP counts at different steps of the SNP discovery pipeline. The percentage of SNPs located on chromosomes or scaffolds remaining after each step are shown in parentheses. The first percentage is relative to the initial set of SNPs and the second percentage is relative to the set of SNPs from the previous step.

### Genotyping the RIL population

Using the per line transcriptomic data, each RIL was genotyped as having either the R500 or IMB211 allele at each of the 149,097 SNPs identified from the population-based SNP-discovery pipeline. An representative RIL genotype plot is shown in Figure 2.

**Figure 2.**
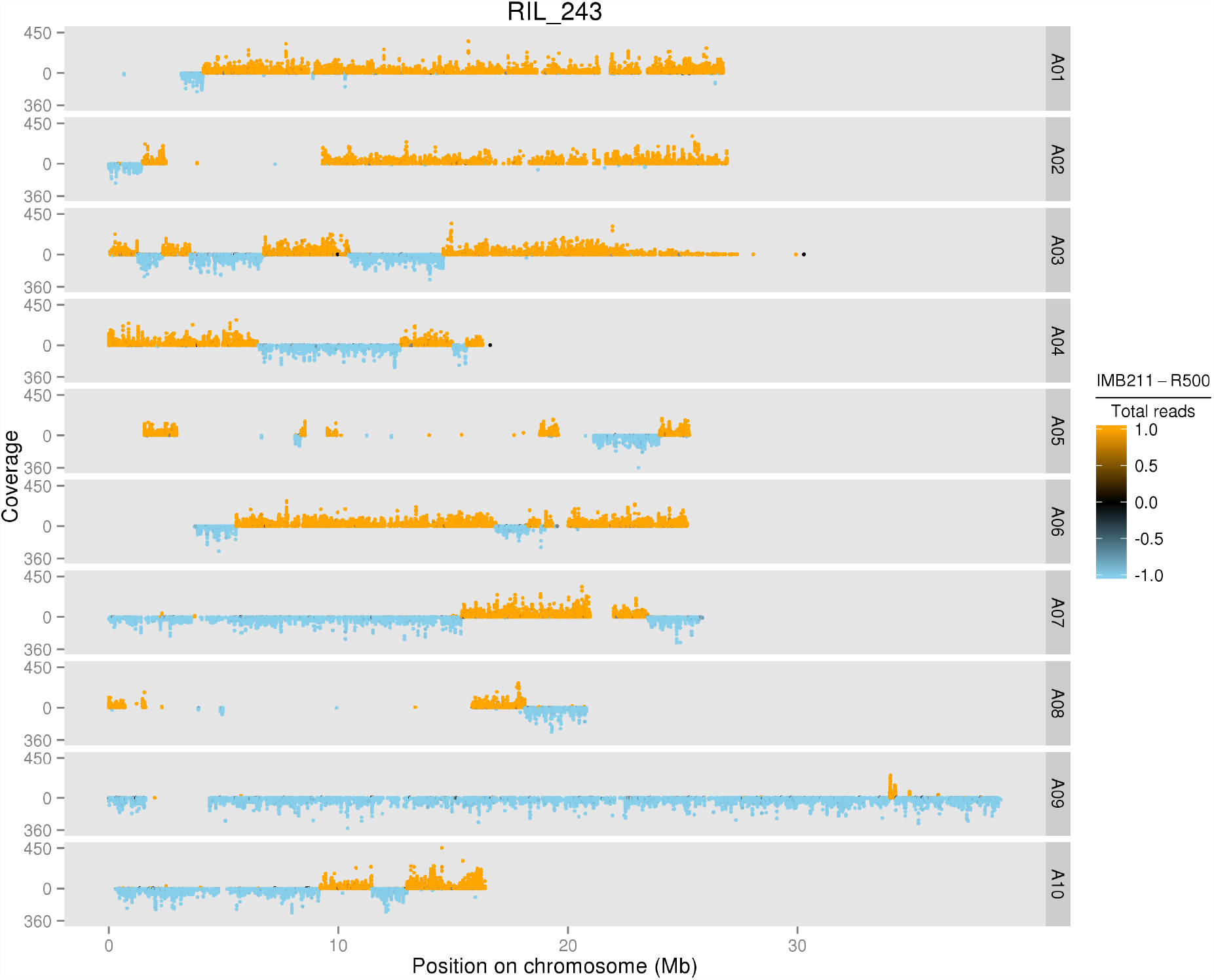
An individual plot of a RIL genotyped with the population based SNP set. Each of the *B. rapa*ten chromosomes are displayed (A01-A10) with counts coverage of each SNP at each physical position on the chromosome in megabases (Mb). The color indicates the relative ratio of coverage between R500 and IMB211 for every SNP.

### Collapsing Adjacent SNPs into Population-Wide Genotype Bins

The next step towards creating a new genetic map was to define the largest set of non-redundant SNPs. This is necessary because the 149,097 SNPs in the full dataset vastly exceed the expected number of recombination breakpoints in a population of 124 individuals. We developed a method to identify and summarize the “genotype bins” in the population. First, we found all detectable recombination breakpoints for each RIL. Next, we consolidated these breakpoints for the entire population. SNPs that were not adjacent to a recombination breakpoint in any of the RILs were considered redundant and removed. This yielded bins of adjacent SNPs with genotype patterns that differed from neighboring bins for at least one RIL because of a recombination event in that specific RIL. The genotype bins for the RIL population are summarized in a composite population genotype map (Figure 3).

**Figure 3.**
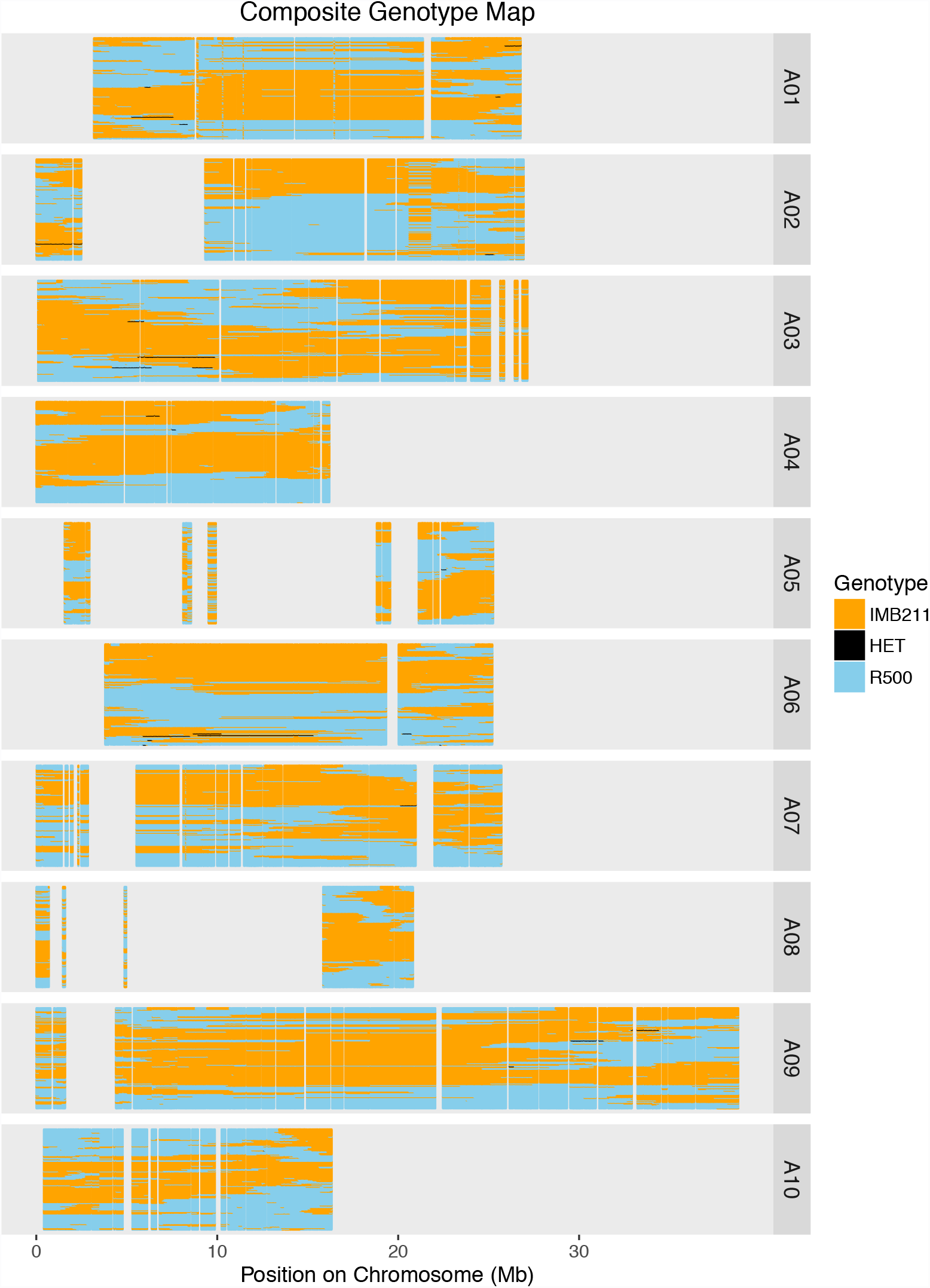
Composite population genotype map with the physical position for each of the ten chromosomes. Each RIL is represented as a single row displaying the genomic region inherited from IMB211 (Orange) or R500 (Blue). Small heterozygous regions are represented in black.

### Finding and Reassigning Misassembled Genomic Regions

A first version of the population genetic map revealed several markers that seemed to be misplaced based on physical position, resulting in large genetic distances between them (Supplemental Figure 2). Given, that we have corrected the parental genotyping issues, the mostly likely explanation for this finding is that these regions represent genome assembly errors. To test this hypothesis, the genotypes of representative SNPs from each bin were used to calculate the asymmetric binary distances between each bin across the population. If the predicted genome position of each bin is correct, the expectation is that each SNP should have the lowest distance to adjacent SNPs in genome coordinates. However, consistent with genome assembly problems, there were a subset of SNPs whose genotypes were more highly correlated with SNPs located elsewhere in the genome rather than with SNPs near their current assigned genomic position (a representative example is shown in Figure 5). To fix these issues, 13 regions consisting of 19 genotypic bins were moved to different genomic locations and 4 regions consisting of 66 bins were inverted in place at their original position based on assymetric binary distance (Supplemental Table 2).

**Figure 5.**
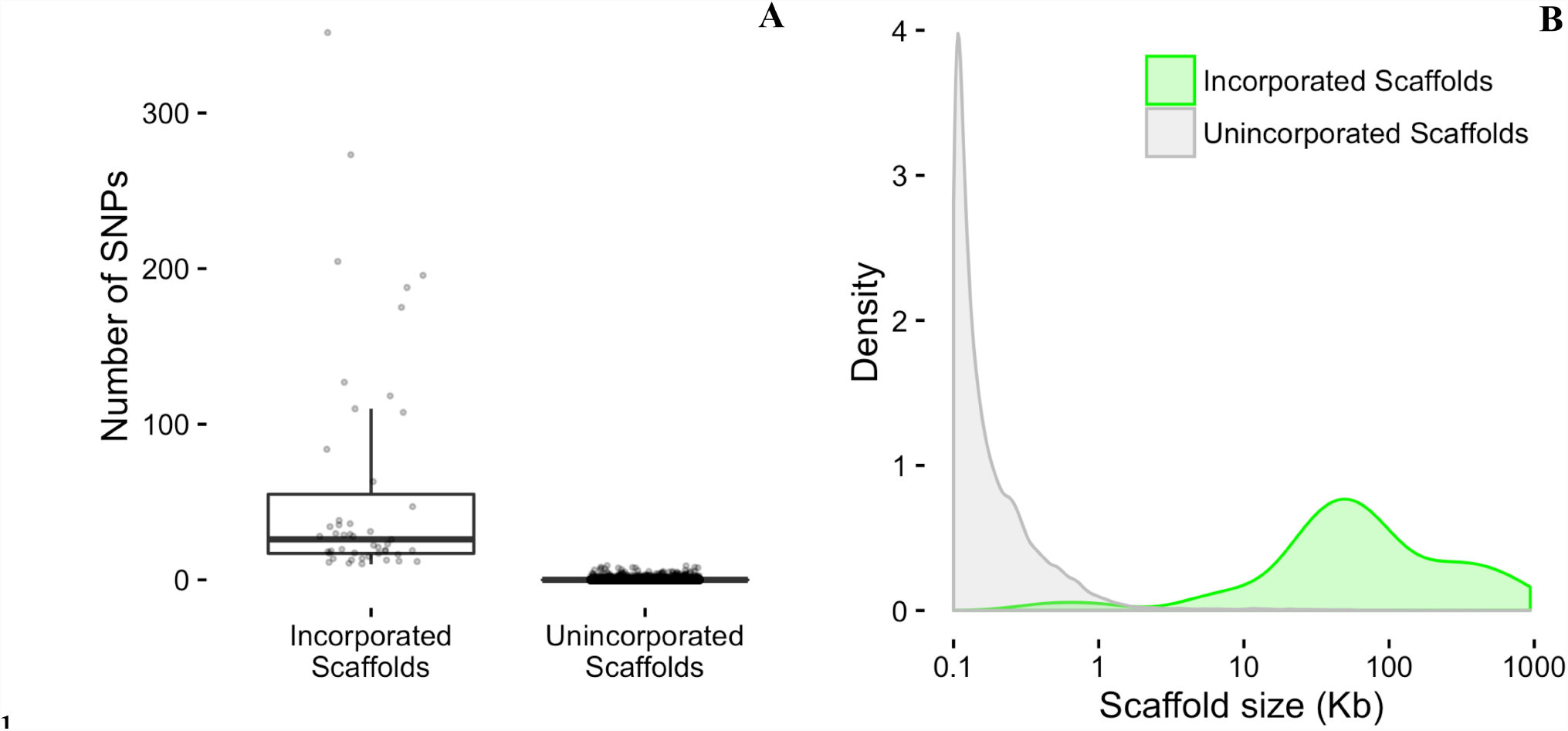
(A) Number of SNPs per scaffold. (B) Density distributions of scaffold sizes. Newly incorporated scaffolds are shown in green and unincorporated scaffolds are shown in gray.

### Incorporating scaffold sequences into the genome

In the current version of the *B. rapa*genome annotation (v1.5) there are 40,357 scaffolds that have not been incorporated into any of the ten chromosomes. These scaffolds range in size from 100 bp to 938 Kbp and represent 1,411 genes spanning 27.5 Mbp. For comparison, there are 39,609 genes within the 283.8 Mbp of annotated chromosomal sequence. Given that the scaffolds contain about as many genes as would be expected on one third of an average chromosome, we decided to extend our strategy for fixing genome misassembles to estimate the approximate chromosomal locations of the scaffolds.

We identified 3,070 SNPs across 339 of the 40,357 scaffolds (the remaining scaffolds had no SNPs, Supplemental Table 3). To be confident in our placement we limited ourselves to the 47 scaffolds with 10 or more SNPs. For each of these 47 scaffolds, we were able to identify at least one genomically defined chromosomal bin that had identical or near identical genotypes. This indicates a very close genetic linkage between the unplaced scaffold and the placed genomic bin allowing us to assign a genomic position to the unplaced scaffold, but not the orientation. The incorporated scaffolds range in size from 429 to 884,746 bp and are enriched for larger scaffolds (Figure 4). The incorporation of these 47 scaffolds allowed us to incorporate 25% (~ 7Mbp) of the unplaced genomic sequence into the genome, representing 49% (691) of the unplaced scaffold genes (Table 2; Supplemental Table 2).

**Figure 4.**
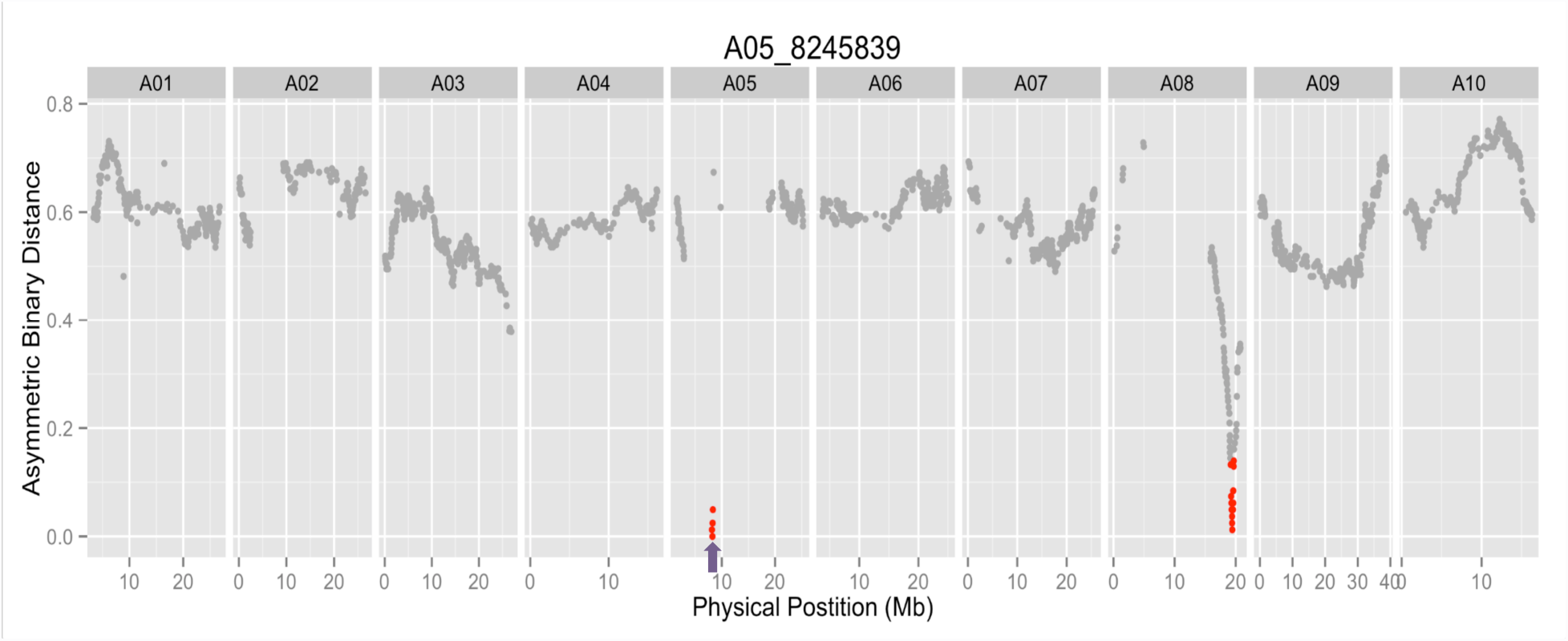
A representative asymmetric binary distance plot for a single molecular marker, A05-8245839, indicated by the purple arrow. Markers with 90% correlation to A05-8245839 are indicated in red and occur on chromosomes A05 and A08. The group of markers on A05 were moved to A08.

**Table 2.** Incorporated scaffolds represent a disproportionately high amount of scaffold sequence. Percentages of scaffold subset counts and total lengths relative to the set of all scaffolds are shown in parentheses.

|  | Count | Total Length (bp) | Mean<br>Length (bp) | Median Length (bp) |
| --- | --- | --- | --- | --- |
| Incorporated<br>Scaffolds | 47 (0.1%) | 6,927,293 (25.1%) | 147,389 | 58,889 |
| Unincorporated<br>Scaffolds | 40,310 (99.9%) | 20,655,028 (74.9%) | 512 | 140 |
| All Scaffolds | 40,357 | 27,582,321 | 683 | 140 |

While most of the incorporated scaffolds represent a single genotype bin, seven scaffolds are comprised of multiple bins. Scaffold000164, for example, includes 65 annotated genes across six distinct genotype bins within its 313.7 Kbp sequence. For six of the scaffolds with multiple bins the bins were closely linked and allowed us to place the scaffold in a single location in the genome. However, one scaffold, Scaffold000191, contained two bins that mapped to two different chromosomes, indicating that it was misassembled. Therefore, we split its two bins and assigned them to the appropriate chromosome locations (5 genes/28.2 Kbp to A01 and 24 genes/104.1 Kbp to A05)

Possible reasons for the enrichment of larger scaffolds within the set of incorporated scaffolds include: (1) larger scaffolds are more likely to include expressed genes and, therefore, SNPs that we can detect and (2) larger scaffolds may be more likely to be accurate representations of a contiguous region within the genome. This second point is based on the assumption that large scaffolds could be assembled perhaps due to more abundant, more consistent, and/or more convincing experimental support than small scaffolds. Before (Figure 6A) and after (Figure 6B) plots of genome-wide asymmetric binary distances for each marker pair show that rearranging putative genomic misassemblies and incorporating scaffolds eliminates inconsistencies between genome position and genotypes of adjacent markers.

**Figure 6.**
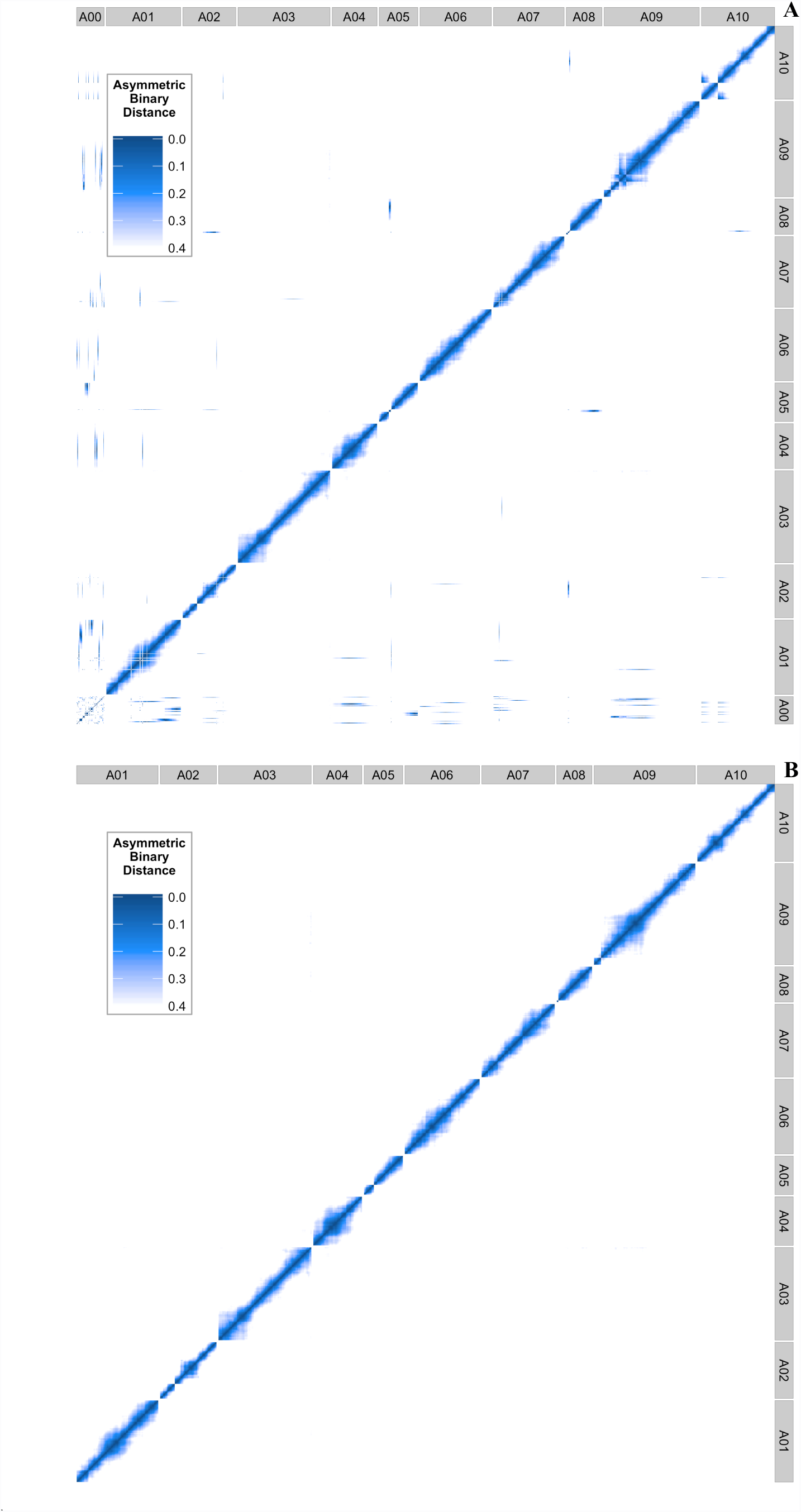
Genome-wide asymmetric binary distance plots for each marker compared against every other marker (A01-A10). A00 in panel A represents the unplaced genomic scaffold sequences. Dark blue indicates high correlation (low asymmetric binary distance), while white indicates no correlation. The right plot is the final position of each marker and scaffold after applying our pipeline.

### High-density genetic map

From the available SNP data we were able to create a genetic map with ten linkage groups corresponding to the 10 chromosomes of *Brassica rapa*. The map contains 1482 genotyped markers for 124 RILs and is effectively saturated based on recombination events existing in the population (Supplemental Table 4). The new map has an average marker spacing of 0.7 cM and a total map distance of 1,045.6 cM. For comparison, the original map contained 225 markers with an average spacing of 3.3 cM (Iniguez-Luy *et al.*, 2009). This is also compared to a recent map created on a subset of the population, 67 RILs, that had a total of 125 markers derived from microarray probes (Hammond et al., 2011). Having the genetic distance of markers with known genomic coordinates allowed us to fix two additional genome misassembles resulting in large inversions on chromosomes A09 and A10 (Figure 7). All of these improvements combined allow us to more accurately map QTL for known phenotypes such as flowering time (Figure 8). Lastly, we fit spline based regressions for each chromosome to more accurately convert between genetic distance and physical distance (A01 example; Figure 7B, C). These conversion equations are helpful for finding candidate genes in significant QTL regions (Fulop *et al.*, 2016).

**Figure 7.**
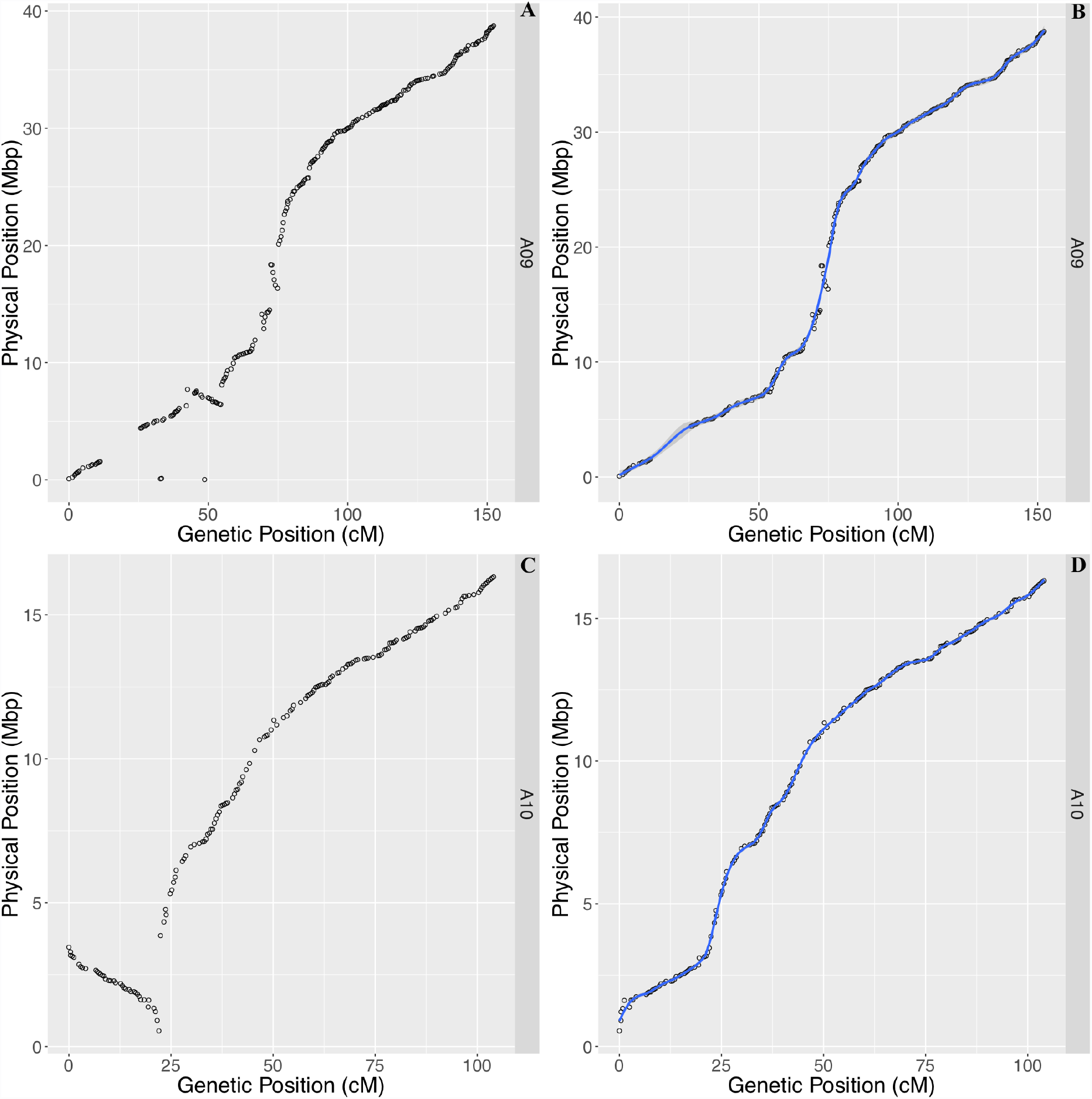
Physical position versus genetic position of each marker for chromosome A09 (A,B) and A10 (C,D) using genome version 1.5 (A,C) and fixed inversions using recombination information (B,D). Loess smoothing for converting between genetic and physical distance is displayed by the blue line in panels B and D.

**Figure 8.**
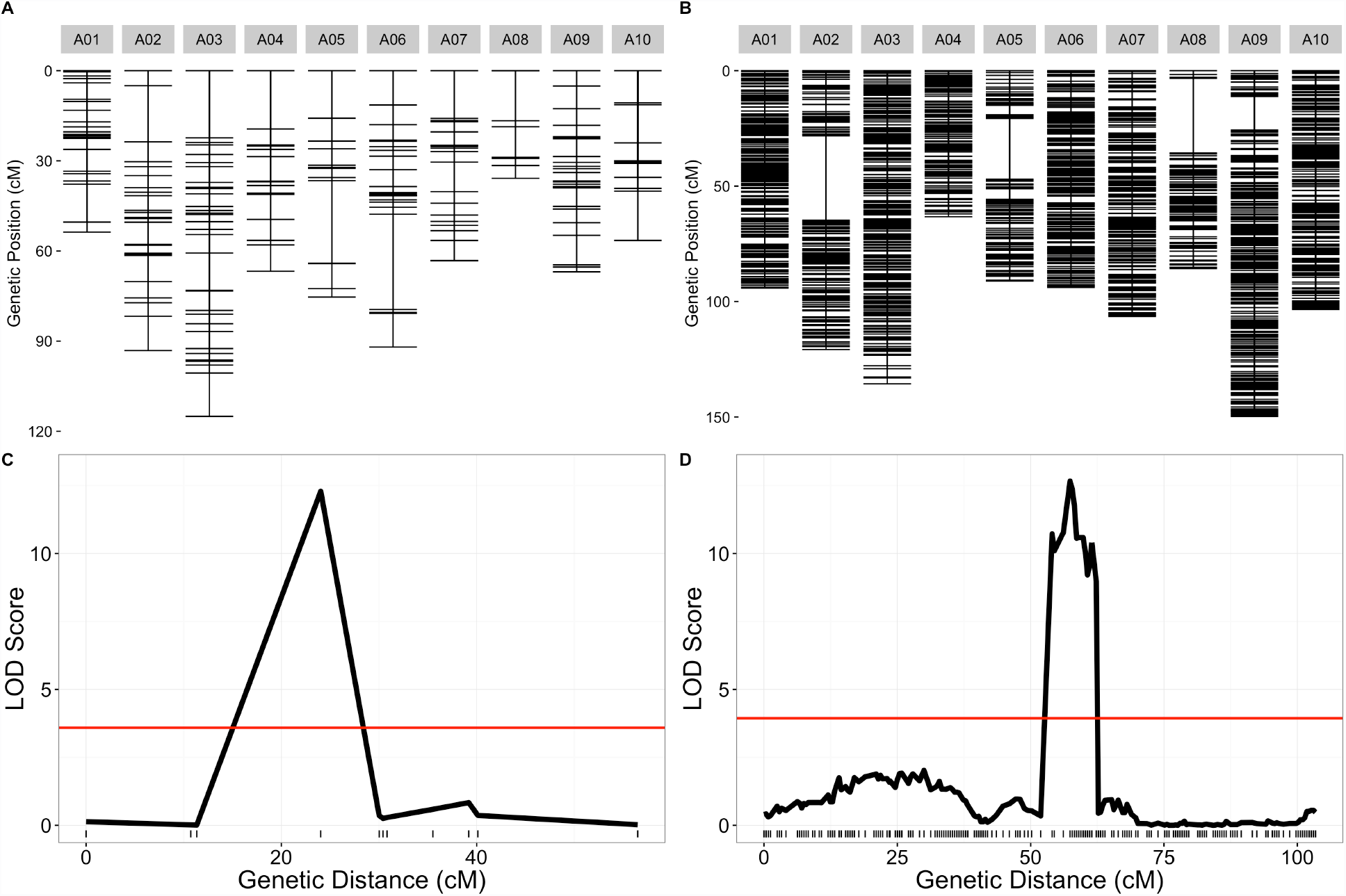
Old and new genetic map comparisons. Genetic markers for each chromosome are displayed in centimorgan distance (cM) for the old (A) and new (B) genetic maps. Comparison of likelihood odds scores for flowering time QTL using the old (C) and new (D) genetic maps.

## CONCLUSIONS

In this study we demonstrated the flexibility and power of thoughtfully designed RNA-seq experiments from tissue collected from a field experiment. RNA is a rich source of biological information that can be utilized beyond expression analysis and transcriptome annotation. It is our hope that these new community resources using RNA-seq are used to further genome annotation, assembly, and functional analysis of the emerging model crop *Brassica rapa*.

## Acknowledgments

The authors wish to thank Maloof Lab members for helpful discussion and reading of the manuscript. RJC Markelz was supported by a NSF Postdoctoral Research Fellowship in Biology (IOS-1402495). This research was supported by NSF grant IOS-0923752 to CW and JNM.

## Supporting Information

Supporting Code for genetic map construction can be found at:https://github.com/rjcmarkelz/brassica_genetic_map_paper

## Supplemental Data

Supplemental Figure 1-Genetic Map before misplaced markers reassigned. Supplemental Table 1-RIL population SNP genotyping and genomic position. Supplemental Table 2-RIL population genetic bins and scaffold original and final positions. Supplemental Table 3-SNP base pair calls on scaffolds. Supplemental Table 4-Final genetic map of RIL population.

